# Large Language Models for Accessible Reporting of Bioinformatics Analyses in Interdisciplinary Contexts

**DOI:** 10.1101/2025.11.09.687479

**Authors:** Lijia Yu, Daniel Kim, Yue Cao, Matthew Wei Shun Shu, Maya Shen, Xiaoqi Liang, Jasmine Gu, Rojashree Jayakumar, Wenze Ding, Fei Yang, Xumou Zhang, Jinman Kim, Pengyi Yang, Jean Yee Hwa Yang

## Abstract

Health and life scientists frequently rely on quantitative experts to perform complex data analyses, yet interpretation of these results often becomes a bottleneck due to communication barriers across disciplines. Large Language Models (LLMs) have the potential to act as intermediaries that translate analytical outputs into accessible scientific narratives, but their ability to reliably interpret outputs from real-world biomedical data analyses across disciplines remains unclear. In this study, we investigate how state-of-the-art LLMs behave when integrated into real bioinformatics analysis workflows as report-generation assistants for interdisciplinary teams. We evaluated LLM-generated summaries using both automated and human evaluation frameworks to ensure holistic evaluation. Automated assessment employed multiple choice questions designed using Bloom’s taxonomy to assess multiple levels of understanding, while human evaluation tasked scientists to score summaries for factual consistency, lack of harmfulness, comprehensiveness, and coherence. All generally produced readable and largely safe summaries, confirming their value for first-pass translation of technical analyses, however frequently misinterpreted visualisations, produced verbose summaries and rarely offered novel insights beyond what was already contained in the analytics. Our findings suggest that LLMs are best suited for easing interdisciplinary communication rather than replacing domain expertise and human oversight remains essential to guarantee accuracy, interpretative depth, and the generation of genuinely novel scientific insights.

## Introduction

The rapid advancements in Large Language Models (LLMs) have led to unprecedented capabilities in broad research areas. Built on the transformer architecture, these models contain billions of parameters and have been trained on a large corpus of text, allowing them to capture intricate linguistic patterns and relationships (Naveed *et al*., 2023). Their ability to handle increasingly complex data and tasks with remarkable accuracy has pushed LLMs as foundational building blocks for both specialized and general-purpose AI agents in diverse areas of research. These include health care, medicine, and particularly bioinformatics (Clusmann *et al*., 2023a; Moor *et al*., 2023; Sarumi and Heider, 2024), where they have been fine-tuned as specialized agents for applications in protein structure predictions, biological sequence analysis, drug discovery, and more (Yang *et al*., 2022; Zhou *et al*., 2023; Sarumi and Heider, 2024; Wong *et al*., 2024). For example, DNABERT-2 and scBERT have adapted the architecture of the NLP model BERT for multi-species genomic analyses and cell type annotation, respectively (Yang *et al*., 2022; Zhou *et al*., 2023). On the other hand, Lomics leverages multiple LLMs to derive relevant pathways tailored to specific scientific research questions in an interpretable output (Wong *et al*., 2024).

LLMs have the potential to expand beyond enhancing biomedical data analysis; they offer unique advantages in parsing complex scientific results into human-interpretable and context-aware formats that substantially improve accessibility (Singh *et al*., 2024). There has been growing attention of LLMs in the context of patient care, where clear communication is a critical determinant of informed decision-making, treatment adherence, and ultimately healthcare outcomes. For example, LLMs show promise in improving communication between healthcare providers and patients via their ability to summarise complex medical information, potentially improving patient comprehension and satisfaction (Ayers *et al*., 2023; Clusmann *et al*., 2023b), and have been deployed to assist in clinical trial recruitment by translating complex medical jargon into more accessible language, reducing barriers to patient participation (Tang *et al*., 2023; Van Veen *et al*., 2024). Recently, an agent system integrated LLMs with databases to automate the discovery of biomarkers and generation of enrichment reports (Pickard *et al*., 2025).

While much of the current emphasis has been on patient-facing communication, similar interpretability challenges exist in scientific research settings. Although LLMs show promise in healthcare and clinical contexts, their potential to synthesise and communicate bioinformatics analyses into concise, interpretable summaries remains largely underexplored. This capability is crucial given the frequent collaboration between biomedical researchers and quantitative scientists, such as bioinformaticians, where communication gaps often arise due to differences in domain knowledge and terminology. Leveraging LLMs offers a promising opportunity to bridge these gaps by generating coherent and context-aware narratives that connect computational insights with biological understanding.

Numerous benchmarks have been developed to evaluate LLMs across a variety of tasks, such as fact recall, mathematical reasoning, and code generation. More recent efforts have pushed into complex workflows. Examples include literature reviews and end-to-end data analyses (Chen *et al*., 2021; Lai *et al*., 2022; Seßler *et al*., 2024; Tang *et al*., 2024) and domain-specific suites like BixBench, which target specialised data-analysis capabilities (Mitchener *et al*., 2025). More recent efforts have extended to complex workflows and systematic evaluations of LLM capabilities (Shang *et al*., 2025; Center for AI Safety, Scale AI and HLE Contributors Consortium, 2026). However, with the exception of a few studies which evaluate the ability of LLMs to interpret different types of data visualisations (Pawelec *et al*., 2024; Hong *et al*., 2025), there remains a gap in benchmarks designed to assess the ability of LLMs to interpret analytical results in context of a research question, particularly within bioinformatics.

To address this need, we evaluated the zero-shot performance of seven state-of-the-art LLMs: including non-reasoning models Gemini 2.0 Flash (publicly available from January 2025), Claude 3.7 Sonnet (February 2025), GPT-4o (May 2024), and reasoning models Gemini 3.6 Flash (July 2026), Claude Opus 5 (July 2026), o1 (December 2024) and GPT-5.6 Sol (July 2026), assessing their ability to perform tasks without having received any training examples. For simplicity we will refer to Gemini 2.0 Flash as Gemini 2.0, Gemini 3.6 Flash as Gemini 3.6, and Claude 3.7 Sonnet as Claude 3.7 from now on. We provided the models with typical bioinformatics analytics from six broad areas: simulation, spatial transcriptomics, pathway analysis, differential expression (DE), classification, and cell-cell interaction (CCI). To comprehensively evaluate the performance of the tested models, we implemented two evaluation tracks: *automated* and *human evaluation tracks*. The *automated evaluation track* included multiple choice questions that were designed using Bloom’s taxonomy (Adams, 2015) and the *human evaluation track* involved domain specialists qualitatively rating each model’s summaries for factual accuracy, absence of harmful content, coherence, and comprehensiveness.

## Results

### Study design for evaluating LLMs in interdisciplinary biomedical collaborations

We designed a study to explore and examine the promise of large language models (LLM) in facilitating the communication of bioinformatics analytic results into interpretable summaries using accessible language, thereby reducing the discipline language barrier between quantitative scientists to health and life science domain experts. We achieve this by developing a collection of bioinformatics analytic reports together with a comprehensive evaluation framework comprising *automatic evaluation* and *human evaluation* tracks to assess the capabilities of seven widely accessible LLMs to translate the analytic reports into coherent summaries: non-reasoning models Gemini 2.0, Claude 3.7, GPT-4o and reasoning models Gemini 3.6, Claude Opus 5, o1, and GPT-5.6 Sol.

The reproducible analytic reports were generated using the R programming language in R Markdown (.Rmd) format and contain text on contextual information, code, data and/or graphical output. For the *automatic evaluation track,* we curated 117 representative analytic reports, from 46 bioinformatics case studies, spanning a range of common bioinformatics analytics (see Supplementary Figure S1). The biological-focused analytics category includes differential expression, pathway analysis, and cell-cell interactions and the computational-focused analytics category includes simulation benchmarking, classification, and spatial analysis. Each case varied in its input type, including different combinations of code, graphs and data (see Supplementary Figure S1). Each of these R-based analytic reports were passed to a LLM (*Report-generator*), which was tasked with interpreting and synthesising the critical information in the analytic report to produce a LLM summary report with the target audience being experimental scientists (Figure 1A). The summary reports output by the *Report-generator* and a set of corresponding multiple choice questions (MCQs) were passed to a second LLM (*MCQ-responder)* which was tasked with answering the MCQs using the information contained in the summary report.

**Figure 1.**
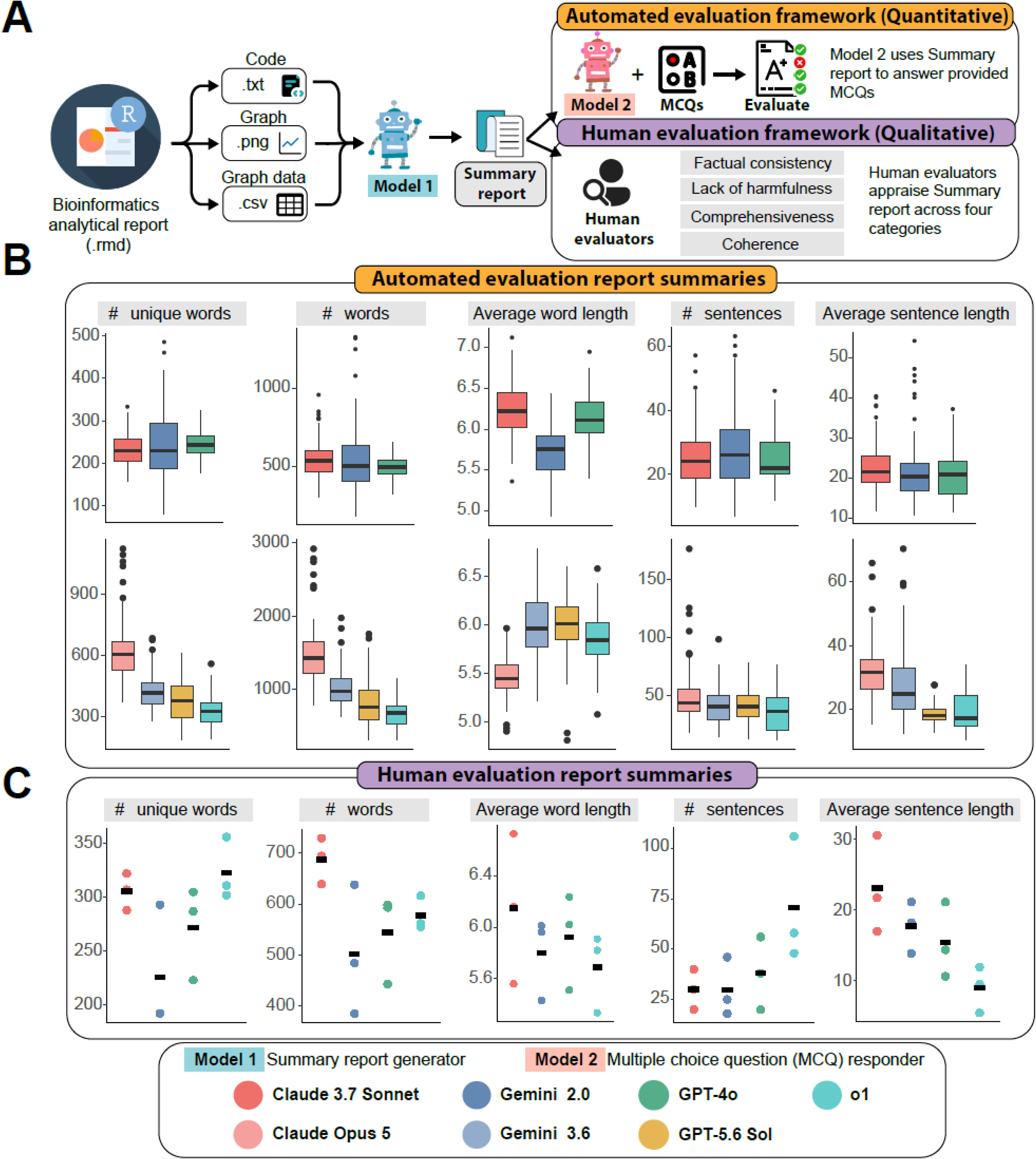
Framework for benchmarking large language models on bioinformatics report generation. (A) Bioinformatics analytic reports containing code and graphs are generated across a collection of analysis topics. The analytic reports are passed into LLMs to generate summary reports. The quality of the summary reports is assessed through automated evaluation and human evaluations. These LLMs are termed summary report generators. In automated evaluation, the summary reports are passed to LLMs to answer multiple-choice questions (MCQs) based on the generated report. These LLMs are termed multiple choice question responder. The responses are compared against reference answers for accuracy. In human evaluation, we selected three representative summary reports and asked expert reviewers to appraise the reports across four criteria. (B) shows the characteristics of the summary reports evaluated in the automatic evaluation. (C) shows the characteristics of the three representative summary reports selected for human evaluation.

For the *human evaluation track,* we curated three extended analytic reports containing all the code, figures, and underlying data in the analytic report across three case studies: pathway analysis, cell-cell interaction, and classification. The three extended analytic reports were passed to the *Report-generator* and the resulting summary reports were then rated for factual consistency, lack of harmfulness, comprehensiveness, and coherence by reviewers from varied disciplinary backgrounds using a structured survey (Figure 1A).

To examine differences in textual characteristics of all summary reports generated by the different LLM models, we performed a one-way ANOVA on all the summary reports generated from the *Report-generators* for both evaluation tracks. Across all the case studies assessed using the *automated evaluation track* (Figure 1B), we observed statistically significant differences within both non-reasoning and reasoning LLM groups in average word length (*F*_2,348_=95.94, *p*<0.01; *F*_3, 464_ = 106.7, *p*<0.01), total word count (*F*_2,348_=7.511, *p*<0.01; *F*_3, 464_ = 175.2, *p*<0.01), and number of sentences (*F*_2,348_=3.459, *p*=0.0325; *F*_3, 464_ = 8.764, *p*<0.01). Among the reasoning models, Claude Opus 5 produced the highest total word count and the greatest number of sentences on average, whereas GPT-5.6 Sol produced the longest words on average. Among the non-reasoning models, no significant differences were observed in the number of unique words (*F*_2,348_=1.43, *p*=0.241) or average sentence length (*F*_2,348_=2.025, *p*=0.134) among non-reasoning models. For the three extended analytic reports assessed using the *human evaluation track* (Figure 1C), we found no stylistic variation between the LLM models (GPT-4o, o1, Gemini 2.0, and Claude 3.7) except average sentence length (*F_3,8_*=4.11, *p*=0.049).

### LLM summaries preserve the majority of analytic content

We assessed each *Report-generator* by measuring how well its summaries recapitulated the key information contained in the analytic reports. Here, we tested all non-reasoning and reasoning LLM groups. Performance was assessed within each group by providing each summary, along with corresponding MCQ, to the *MCQ-responders* and calculating the average accuracy. To our surprise, overall performance was lower than expected given the promise of LLMs commonly believed. The top performance in non-reasoning models, an average of 0.61, was obtained from summaries produced by Gemini 2.0 (Figure 2A). Summaries from Claude 3.7 and GPT-4o followed with 0.56 and 0.48, respectively. The reasoning models showed consistently higher performance, with Gemini 3.6 achieving the highest mean accuracy overall (0.75), followed by Claude Opus 5 (0.68), GPT-5.6 Sol (0.67), and o1 (0.65). We performed a sensitivity analysis (Figure 2A), which demonstrates the accuracies are similar regardless of which LLM model was the *MCQ-responder*, indicating that the ability to answer the MCQs are due to the quality of the generated summaries themselves, and not affected by the choice of *MCQ-responders*.

**Figure 2:**
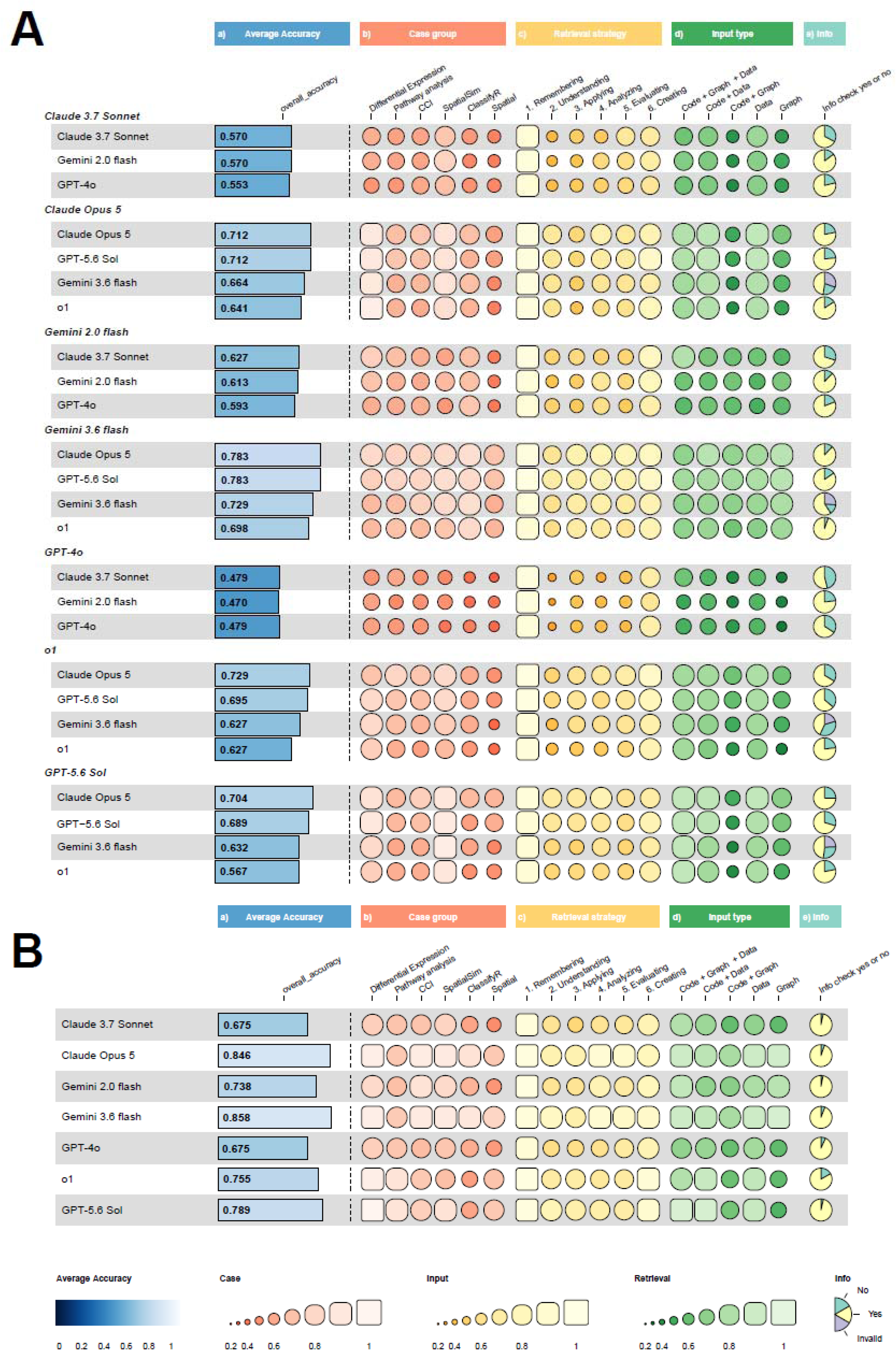
Overview of the automatic evaluation result. (A) shows MCQ accuracy for automatic evaluation and (B) shows MCQ accuracy for baseline. In both panels, column (a) shows each model’s overall average accuracy. Columns (b)-(d) show the rank of accuracy across three dimensions: (b) Case group (e.g., Differential Expression, Pathway Analysis, Cell-Cell Interaction), (c) Retrieval strategy (MCQ design mapped to cognitive levels from Remembering to Creating), and (d) Input type (e.g., Code, Graph, Data, or combinations). Column (e) shows whether the model reports identifying relevant information (“Info check”), represented as pie charts.

The error in MCQs may stem from two sources: (1) the report-generating LLM fails to pass along critical information during its summarisation, or (2) the MCQs are too challenging for the LLM to solve. To disentangle the two and better quantify the LLM’s capacity in summarising information, we should first establish a baseline performance metric to measure the difficulty level associated with each MCQ question. We established the “baseline” performance by having each LLM model to answer the same MCQs directly from the original analytic report with data and graphs. Next, we defined relative accuracy as the ratio between the “summary report-based accuracy” and “baseline” performance. We believe this is a more appropriate metric as it also accounts for “baseline” differences between different cases. Using this more meaningful metric, we observed that summary reports generated by the non-reasoning models Claude 3.7 and Gemini 2.0 corresponded to a relative accuracy of 83-84%, while GPT-4o achieved 70% of its optimal baseline (Figure 2B). Among the reasoning models, Gemini 3.6 achieved the highest relative accuracy at 87%, followed by o1 (86%), GPT-5.6 Sol (85%), and Claude Opus 5 (81%).

Based on these observations, we hypothesised that MCQ performance depends strongly on whether the information required to answer each question is preserved in the generated summary. For each MCQ, we asked the question, “Does the analytic or summary report contain the information necessary to answer the provided MCQ?” The information coverage metric was then calculated as the proportion of “Yes” responses. In the baseline case as defined in the previous paragraph, information coverage was consistently high across models, ranging from 83.2% to 96.9% , indicating the relevant information is indeed contained in original analytic reports. In contrast, the information coverage declined in the summary reports generated by the *Report-generator*. Among the non-reasoning models, Gemini 2.0 retained the highest information coverage (79.6%), followed by Claude 3.7 (77.0%) and GPT-4o (65.4%). Among the reasoning models, Gemini 3.6 achieved the highest information coverage (87.3%), followed by Claude Opus 5 (77.5%), GPT-5.6 Sol (72.2%), and o1 (65.8%). These results suggest that the use of reasoning models did not uniformly improve information retention during report summarisation (Supplementary Figure S2). We next examined whether the availability of the required information was associated with MCQ accuracy. As expected, when the required information was present in both the original analytic report and the generated summary, all seven LLMs achieved high absolute accuracy (0.65-0.81), corresponding to 83%-97% of their respective baseline accuracy. In contrast, when the information was present in the original analytic report but absent from the generated summary, absolute accuracy decreased substantially to 0.15-0.45, corresponding to only 26%-56% of baseline performance (Figure 3A). Together, these findings indicate the importance of extraction of relevant information in the report and the need of strategies such as customised prompting or guided reporting to ensure essential analytical content is preserved.

**Figure 3:**
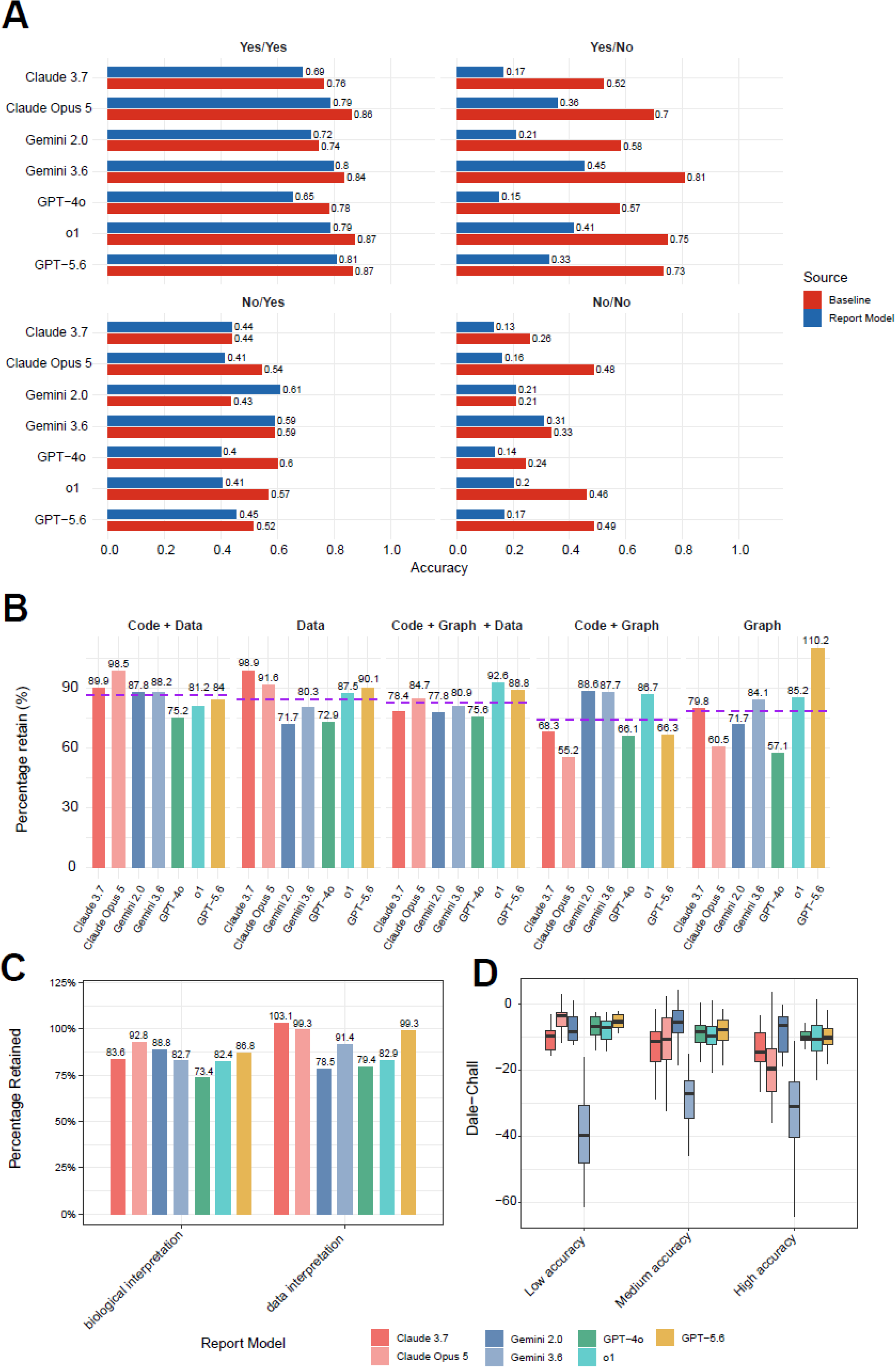
Performance comparison on automatic evaluation accuracy. (A) Accuracy of baseline model versus report models for each report-generating LLM. “Yes” and “No” denote whether the information is available in the report under each scenario. For example, “Yes/No” indicates the information is available in the baseline model but not in the report models. (B) Information retention across different input modalities for each report-generating LLM. Information retention is calculated as the ratio between information availability in the report model and in the baseline model. Purple dashed line indicates the average across all LLMs. (C) Information retention across report models stratified into biological focused and data focused case studies. (D) Readability of generated reports measured using the Dale–Chall readability index, average MCQ accuracy was computed and stratified into three categories: low (<0.3), medium (0.3-0.7), and high (>=0.7).

### Input type influences information retention in LLM-generated reports

We took a closer look at whether the amount of information captured during the generation of the summary report is affected by factors such as input type and analysis type. In terms of input type, we compared using data exclusively, graph exclusively, code + data, code + graph, and code + graph + data as input to the *Report-generator*. On average across the seven models, the *Report-generators* generated summary reports retained more information when the analytic reports contained data compared to graphs (Figure 3B). Specifically, code + data achieved a high relative accuracy of 86.4%, followed closely when the input is data only with relative accuracy of 84.7%. In comparison, graph only and code + graph achieved lower mean information retention of 78.4% and 74.1%, respectively. An exception to this overall trend was GPT-5.6 Sol for graph-only inputs, where information retention reached 110.2% relative to baseline. This indicates that the performance based on the generated summary slightly exceeded that obtained directly from the original input. These results highlight the current challenge of interpretation using graphs, which could introduce more opportunities for error than directly reading it from data or code for LLM, consistent with current limitations of LLM models (Zhang *et al*., 2025). When looking at the type of MCQs, we noted that code + graph is especially weaker at the Level 2 (Understanding) questions (Supplementary Figure S3). Questions at this level involve straightforward interpretations of the data, such as determining the number of patients in each disease group or the total number of cell types, where reading these basic statistics directly from the data may produce more accurate interpretations compared to inferring them from a graph alone.

To evaluate whether information retention differs by analysis type, we compared MCQ accuracy on summary reports generated from biological-focused and computational-focused analytic tasks (see Methods). Notably, we observed variation in relative accuracy across both models and analysis types (Figure 3C). When Claude 3.7 was used as the *Report-generator*, MCQ accuracy was higher for data interpretation tasks (103.1% relative accuracy) compared with biological interpretation tasks (83.8% relative accuracy), with Claude Opus 5, Gemini 3.6, GPT-4o, and GPT-5.6 Sol showing similar patterns (99.3% vs 92.8%, 91.4% vs 82.7%, 79.4% vs 73.4%, and 99.3% vs 86.8%, respectively). Gemini 2.0 and o1 showed the opposite trend, achieving higher relative accuracy for biological interpretation tasks (88.8% and 82.4%, respectively) than for data interpretation tasks (78.5% and 82.0%, respectively). These results suggest that the effectiveness of a given LLM may depend on the type of analysis, with certain models better suited for interpretation of computational outcomes and others suited for interpreting biological information.

We next stratified MCQ accuracy of the summary reports by Bloom’s taxonomy to assess whether certain types of cognitive tasks were better supported by LLM-generated summaries. We observed two notable extremes: questions at Level 1 (Remembering) and Level 6 (Creating) both resulted in close to 100% relative accuracy (Figure 2A, Supplementary Figure S4). In contrast, questions at the intermediate levels (Levels 2-5) yielded 70-80% relative accuracies. We speculate that this pattern arises because questions at Levels 1 and 6 generally do not require LLMs to interpret the derived data or perform any computational calculation. Level 1 questions typically involve simple fact recall and thus rely primarily on whether the LLM accurately included basic contextual information. Level 6 questions generally required synthesis of information in the report, such as linking enriched pathways or GO terms to broader biological processes. We note that these categories are based on Bloom’s taxonomy and reflect educational cognitive processes within our MCQ framework, rather than true scientific creation or de novo hypothesis generation. Therefore, the high Level 6 accuracy should be interpreted cautiously, as these questions may partly align with LLM strengths in semantic synthesis and biological knowledge retrieval. While seemingly complex, these questions play to the strength of LLMs where LLM are able to leverage its extensive knowledge base.

### Readability of summary report varies across LLMs

Next, to assess whether performance was influenced by the readability of the summary reports in the *automated evaluation track*, we examined word complexity using the New Dale-Chall score (Chall and Dale, 1995) and ease of understanding using the Flesch-Kincaid score (Flesch, 1948). Claude 3.7 and Claude Opus 5 produced summary reports with more complex vocabulary (Dale-Chall: -13.2 and -14.8, respectively), while Gemini 3.6 showed the most extreme score (−32.2). For Flesch-Kincaid scores, a readability metric, Claude Opus 5 produced the most difficult-to-read text (19.2), followed by Gemini 3.6 (18.9) and Claude 3.7 (18.3). Gemini 3.6 and Claude Opus 5 showed the strongest negative Pearson correlations between the two readability scores (*r* = -0.830 and -0.814, respectively), followed by GPT-4o (*r* = -0.700). More moderate negative correlations were observed for Gemini 2.0, Claude 3.7, and o1 (*r* = -0.619, -0.551, and -0.413, respectively), while GPT-5.6 Sol showed no significant correlation (*r* = -0.119, *p* = 0.203) (Supplementary Figure S5). Across all *Report-generators*, there is a significant negative correlation between New Dale-Chall readability scores and MCQ accuracy (cor = -0.27, p <0.001), indicating that more difficult or technically detailed reports (i.e., those with lower readability scores) tend to be associated with higher MCQ accuracy (Figure 3D). Notably, Gemini 3.6 exhibited substantially lower New Dale-Chall scores than the other models across all accuracy groups, suggesting that its generated reports contained a higher density of complex vocabulary.

In the *human evaluation track*, we explored whether readability of the summary reports from the three extended case studies were associated with higher scores by the human evaluators. To do this, we calculated lexical and readability features for each extended report in the *human evaluation track* and, within each questionnaire category (*Coherence, Comprehensiveness, Factual consistency, Lack of harmfulness*), applied the Jonckheere-Terpstra test (Supplementary Figure S6). We found a statistically significant positive association between average sentence length and evaluator scores for the *Lack of harmfulness* category (JT=458.5, *p* =0.04). Hence, evaluators scored reports with longer sentences as making less harmful claims while the number of sentences was negatively associated with evaluator scores (JT=185.5, *p* =0.004), indicating that reports with more sentences tended to receive lower ratings for the *Lack of harmfulness category*. We observed the same negative trend for the number of unique tokens used in the report and *Lack of Harmfulness* (JT=175.5, *p* =0.006) and *Comprehensiveness* (JT=535, *p* =0.03).

To examine associations between syntactic complexity and the evaluator’s scores, we also examined associations between the Flesch-Kincaid Grade-Level and evaluator scores for all four categories across the *Report-generators*. While positive associations were observed across all question categories, only the association for *Coherence* category was statistically significant (JT=1825, *p* =0.01). This suggests that reports with greater syntactic and lexical complexity were perceived as more coherent potentially due to their clearer logical structure, use of cohesive devices, and stronger flow of ideas typical of scientific journal articles.

### LLMs differ in balancing scientific style, accuracy, and depth of interpretation

To evaluate the quality of summary reports via manual evaluation, we tasked human evaluators to score the reports across the four aforementioned questionnaire categories (*Coherence, Comprehensiveness, Factual consistency, Lack of harmfulness)* (Figure 4A, B). To capture which categories the models performed stably and variably in, we calculated Shannon’s entropy of the distribution of evaluator scores for each category (Figure 4C). While *Factual Consistency* and *Comprehensiveness* showed greater variability across models (H=2.43 and H=2.16, respectively), performance for *Lack of Harmfulness* and *Coherence* was generally stronger and less variable (H=1.74 and H=1.79, respectively). Across all models, although they produced non-offensive, easy-to-read reports, they remained largely superficial and rarely extended beyond what was already available in the original analytics report. In addition, we explored lexical similarities between the reports using a Bag-of-Words representation followed by dimensionality reduction, but did not identify any informative clusters, suggesting the differences between the reports for the categories were not confounded (Supplementary Figure S7).

**Figure 4.**
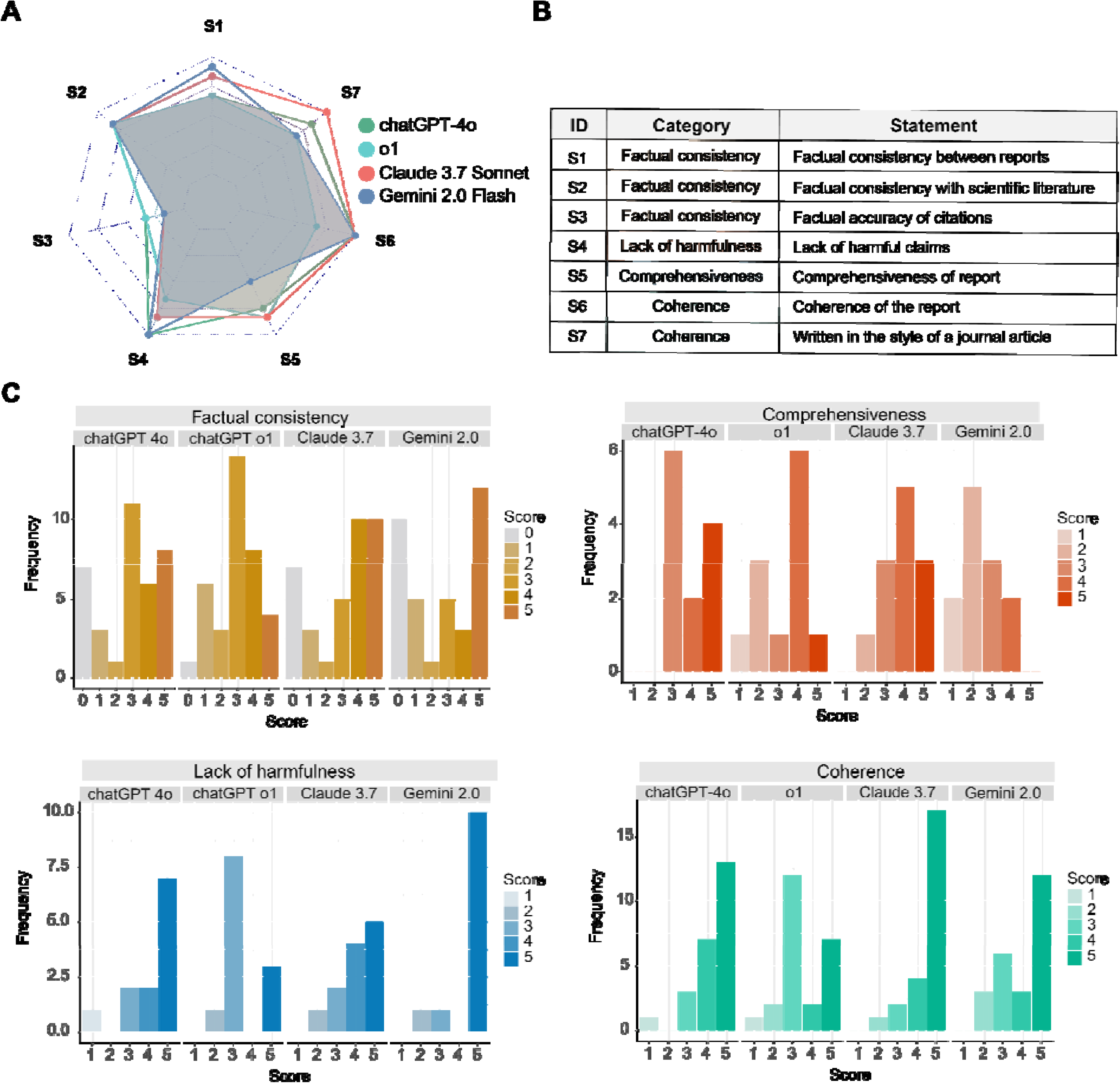
Human evaluation results. (A) Average evaluator scores for each statement in the human evaluation questionnaire across categories. (B) Table of the statements and their corresponding IDs and category assignment in the human evaluation questionnaire. (C) Distribution of evaluator-assigned scores across four evaluation categories: *Factual consistency*, *Lack of harmfulness*, *Comprehensiveness*, and *Coherence*. Scores range from 0 to 5, with higher values indicating better performance. Each distribution reflects the aggregated evaluations of model-generated “Results” sections from evaluating the analytic reports across three case studies.

Claude 3.7 consistently received the highest scores across all categories. Notably, Claude 3.7 was most effective at summarising analytic results in the style of a scientific journal article, the central task of this study. This trend was held when we stratified results by case study type (CCI, Classification, and Pathway), with Claude 3.7 scoring consistently well across all scenarios (Supplementary Figure S8).

Taken together, these findings suggest that while LLMs can generate accessible summaries of complex bioinformatics output, increased verbosity may raise the risk of factual inaccuracies or harmful claims. Moreover, the tested LLMs appear to be less reliable for deeper analytical and biological interpretation.

## Discussion

Synthesising complex bioinformatics analytics into interpretable, faithful, and coherent description is critical to effective cross-disciplinary collaboration between computational and healthcare or life-science professionals. Language models present a unique opportunity to bridge the present gap and thus, it is critical to assess their multi-modal interpretation capability on representative, real-world research questions and outputs. In this study, we investigated the extent to which LLMs can be reliably used to generate interpretable summary reports from complex bioinformatics analyses, with the aim of bridging the disciplinary gap between quantitative and qualitative scientists. To assess if critical information from analyses are retained in the summary reports, we introduced an evaluation framework that compares publicly available models of GPT-4o, o1, GPT-5.6 Sol, Claude 3.7, Claude Opus 5, Gemini 2.0 and Gemini 3.6 using different combinations of code, text, and graph visualisation as input modalities and through both automatic quantitative and manual qualitative evaluation strategies.

We noted performance not only varied according to the model but also with the information provided as input. Consistent with previous benchmarking studies (Vatsal and Dubey, 2024), we found that prompt engineering for both user and system prompts was important to achieve the desired model outputs for evaluation (Supplementary file S9). Specifically, we found that the models required clear and detailed instructions when tasked with summarising analytic reports into a results section typical of a journal article. For example, during the development of both evaluation strategies, we observed that the models are unable to include quantitative evidence to support their claims unless explicitly instructed to do so in the user prompts. Interestingly, providing additional context, such as informing the model that the intended users were bioinformaticians lacking “the biological knowledge to interpret the results and build a cohesive narrative,” appeared to qualitatively improve the relevance and coherence of the generated summaries (Supplementary file S9). These findings demonstrate that providing LLMs with both clear task instructions and contextual information enhances the quality of their outputs and should be considered carefully when using LLMs.

Our case studies included scatter plots, heatmaps, networks, box plots, and dot plots. When provided with visualisations alone, most MCQ-responders struggled. This aligns with prior work surveying GPT-4o, Claude 3.7 in their ability to interpret charts, which universally performed worse on reasoning compared to fact retrieval tasks (Wang *et al*., 2024). Consistent with known weaknesses of earlier visual language models, we found that the newer multimodal large language models (MLLMs) GPT-4o and Claude 3.7 still struggled with tasks which require parsing compositional elements such as axes titles, reading tick marks or understanding figure legends – tasks classified as understanding in our taxonomy (Yang *et al*., 2023; Mukhopadhyay *et al*., 2024). Although recent reasoning models are designed to provide stronger inference capabilities, our results revealed variable performance in comparison to their non-reasoning counterparts. Poor performance could likely be because of the unique charts specific to bioinformatics that are not well represented during training (Huang *et al*., 2024; Wang *et al*., 2024).

In a recent benchmark study of MLLMs applied to full bioinformatics pipelines, the authors noted that explicit prompting to avoid graph generation led to higher performance on MCQ and open answers, postulating this could indicate a reliance on the raw data (Mitchener *et al*., 2025). Findings from another evaluation study found that models likely defaulted to existing knowledge rather than extracting information from the provided charts (Hong *et al*., 2025). Given LLMs perform poorly when provided charts alone, the models are likely inferring compositional details such as axis variables and output categories directly from the raw data (Meng *et al*., 2024). Thus, the tested models perform better when provided with the code and underlying data behind the graph, compared with the code and graph themselves.

LLMs have demonstrated promising capabilities across a range of tasks, including natural language translation, code generation, summarisation, and general-purpose reasoning. While they show growing proficiency in generating readable and non-harmful summaries of analytic reports, our findings highlight a persistent challenge: their limited ability to provide information beyond what is already contained in the input analytics. This suggests that, despite their fluency, many current models may still lack a deeper capacity for synthesis, interpretation, or abstraction which are crucial for analytic reasoning. The preference for Claude 3.7 by the human evaluators, despite its occasional factual inaccuracies, demonstrates the value evaluators place on interpretive insight over factual recapitulation. In addition, the association between fewer, longer sentences, and higher ratings by human evaluators may reflect an implicit preference toward writing styles that appear more coherent or concise, while they may not necessarily be more factually accurate. As LLMs are increasingly deployed in analytical workflows, there is a need to move beyond just considering metrics such as factual consistency and readability, and to develop frameworks that reward models for their ability to add interpretive value, contextual relevance, and domain-appropriate nuance.

Common benchmarks of LLMs traditionally associated with the biomedical field such as PubMedQA, MedMCQA, BioASQ do not effectively evaluate the combination of statistics and clinical data that bioinformatics typically involves (Jin *et al*., 2019; Pal, Umapathi and Sankarasubbu, 2022; Krithara *et al*., 2023). Furthermore, they do not reflect the typical workflow of practitioners which requires synthesis of insights drawn from visualisation and data analysis. Whilst there exists limited studies that evaluate automated analytical pipelines and chart generation, it is still unclear how reliable LLMs models are for use in the interpretation stage of the bioinformatics workflow post-analysis (Mitchener *et al*., 2025). Consistent with previous research, we observed differences between the automated and human evaluation frameworks (Abeysinghe and Circi, 2024).

This study has several limitations. It evaluates properties that are relevant to interdisciplinary communication, but it does not directly test whether LLM-generated reports improve a reader’s understanding or improve collaborative decision-making. In the automatic evaluation, the original analytical report baseline helps distinguish information loss during summarisation from MCQ difficulty or MCQ-responder limitations, but cannot fully attribute every source of error, and information coverage was not manually validated for all reports. Repeated-run sensitivity analyses showed generally reproducible results, with mean pairwise correlations of 0.88-0.92 for most report characteristics and 62.5-79.2% of MCQs receiving identical answers across five runs. (Supplementary Figure S10). However, this was not extended to all case studies and models due to computational cost. In the human evaluation, we did not include reports written by human domain experts as a reference, as the study was designed primarily to compare models. Future work should address these limitations and directly examine how LLM-generated interpretations are used by researchers from different disciplinary backgrounds. Based on the present results, LLMs appear most appropriate as first-pass assistants for translating bioinformatics outputs into accessible scientific reports. Their outputs still require domain expertise to verify the analysis and provide the interpretive depth needed to draw new scientific conclusions.

## Conclusion

Our study was the first to explore the capabilities of seven state-of-the-art LLMs to interpret typical bioinformatics analyses and generate interpretable summary reports. To achieve this, we developed two evaluation frameworks (automated and human evaluation frameworks) that included a diverse set of case studies and MCQ, with varying input combinations. We found that Claude 3.7 performed well in interpreting the provided bioinformatics analyses, while Gemini 2.0 was notably accurate in recall tasks. Their performance was influenced by the detail and quality of the prompts, demonstrating the importance of prompt engineering for optimal performance. In conclusion, our results suggest that the strengths of the tested LLMs lie in bridging the disciplinary boundaries by translating bioinformatics analyses and other technical results into plain, interpretable language. However, our results also highlight the importance of domain expertise to ensure accuracy, depth of interpretations, and the generation of novel scientific insights.

Our finding suggests that with careful prompt engineering, LLMs hold the potential to break the communication challenges between disciplines by translating bioinformatics analyses and other technical results into plain, interpretable language. However, our results underscore the importance of deep discipline expertise required to produce the in-depth interpretations required for new scientific insights that LLMs are not yet able to provide.

## Material & Methods

### Case study design and input preparation

We designed 46 distinct case studies spanning six key areas of bioinformatics analysis: differential expression (DE), pathway analysis (pathway), cell-cell communication analysis (CCI), simulation benchmarking (SpatialSim), patient outcome classification (ClassifyR), and spatial omics exploration (Spatial). Each case study (identified by CaseStudy_ID in the CaseStudies sheet) includes multiple types of input data (Sample_ID), such as analytical scripts, output tables, and visualisations. In total, we curate 117 analytic reports from the 46 case studies, each represented by an R Markdown-based analysis file, using different types of input data. Each R Markdown file is self-contained and structured to address a specific analytical question. It includes a task description, annotated code, and 1-2 output tables or plots, with the total length limited to approximately one page.

#### [A] Data sources and analytical focus

We categorised the six key areas described in the previous paragraphs into biological-focused and computational-focused tasks. Biological-focused tasks require more interpretation of the biology, such as interpreting lists of differential genes and biological processes. Computational-focused tasks focus more on the understanding of data outputs and computational method outputs.

**Datasets:**

**[i] Kidney dataset:** the raw RNA-seq data and metadata containing patient information are downloaded from the National Center for Biotechnology Information (NCBI) Gene Expression Omnibus (GEO) under the accession number GSE46474, including 20 patients with acute rejection and 20 controls(Günther *et al*., 2014).

**[ii] COVID-19 dataset:** the normalized transcriptional count matrix and metadata are downloaded from Figshare: https://doi.org/10.6084/m9.figshare.12436517. The data are subsetted into 13 moderate samples and 12 critical samples of single-cell RNA-sequencing data of airway immune cells(Chua *et al*., 2020).

**[iii] SpatialSim datasets:** benchmarking result tables for 13 methods across all metrics, obtained from Figure 4 of the published single-cell spatial benchmarking study by Liang et al.(Liang, Cao and Hwa Yang, 2024).

**[iv] Breast cancer simulation dataset:** bulk gene expression data and histologically predicted gene expression data from the TCGA breast cancer cohort, including 54 patients with two ER+/PR+ subtypes, obtained from the GHIST paper(Fu *et al*., 2025).

**[v] METABRIC dataset:** it is a selected subset of the METABRIC breast cancer cohort consisting of 165 samples with no detected lymph node metastasis(Rueda *et al*., 2019). Quantile-normalised mRNA microarray data measured by Illumina HumanHT-12 v3 BeadChip was obtained from cBioPortal by the file named data_mrna_illumina_microarray.txt from the METABRIC data set summary.

**[vi] Breast cancer spatial dataset:** Xenium FFPE Human Breast with Custom Add-on Panel dataset was downloaded from (https://www.10xgenomics.com/datasets/xenium-ffpe-human-breast-with-custom-add-on-panel-1-standard) and downsampled to 7,353 cells.

**[vii] Lung cancer spatial dataset:** Xenium Human Lung Cancer dataset was downloaded from https://www.10xgenomics.com/datasets/xenium-human-lung-cancer-post-xenium-technote and downsampled to 2,567 cells.

**The biological-focused tasks include:**

[i] **Differential expression analysis (DE):** Datasets [i] and [ii] were used. The analysis focused on the biological interpretation of differentially expressed genes (DEGs), which were defined using *limma*. This analysis requires LLMs to summarise key molecular changes and their functional implications across disease conditions.

[ii] **Pathway analysis (Pathway):** Datasets [i] and [ii] were used. The analysis focused on pathway enrichment using GSEA. Ranked gene lists were obtained from the log fold-change values of the differential expression analysis. GSEA was performed with the *clusterProfiler* package. This analysis requires LLMs to summarise key pathways in relation to their biological functions, as well as the patient groups in which these pathways are enriched across different conditions.

[iii] **Cell-cell interaction analysis (CCI):** Dataset [ii] was used to derive cell-cell communications using the *CellChat* package. The analysis focused on cell–cell communication, specifically the signaling interactions between cell types, and contrasted interaction strengths across groups. This analysis requires LLMs to interpret cell type interactions and enriched pathways across disease conditions.

**The computational-focused tasks include:**

**[i] Simulation benchmarking result analysis (SpatialSim):** Dataset [iii] was used. The analysis focused on results visualization and comparison. This analysis requires LLMs to summarise the comparison outcome from figures or tables.

**[ii] Classification analysis (ClassifyR):** Datasets [iv] and [v] were used. The analysis focused on comparing the sample-specific accuracy of breast cancer subtype classification across different types of data. This analysis requires LLMs to interpret comparative classification accuracy across sample groups.

**[iii] Spatial analysis (Spatial):** Datasets [vi] and [vii] were used. The analysis focused on visualizing how cell types are organized spatially, comparing domains identified by clustering with manually annotated regions, and assessing cell–cell communication across different cell types in the spatial data. The analysis requires LLMs to interpret the outputs of the spatial analysis, with a particular focus on numerical comparisons.

#### [B] Preparation of input data for summary reports

Each R Markdown file was designed to simulate a realistic analytical workflow. It included: (1) A task overview summarizing the analysis objective and key background. (2) Annotated code chunks for data processing and visualization. (3) Output in the form of CSV tables and publication-ready figures (PNG format).

All tables included descriptive column and row names, and tables were either derived directly from the analysis code or served as inputs for figure generation. Figures were resized as needed to comply with input size requirements of commercial LLM APIs.

#### [C] MCQ design

For each case study, we developed a minimum of three MCQs grounded in the provided outputs, resulting in 138 MCQs. These questions enable a quantitative assessment of whether LLMs can comprehend and interpret the model-generated reports accurately.

To ensure the MCQs cover a diverse range of difficulty, we designed the MCQs according to Bloom’s taxonomy(Adams, 2015) as described in Table 1. Bloom’s taxonomy is an education framework developed in 1956 to help educators structure learning outcomes. In Bloom’s taxonomy, questions can be classified into one of the six categories: Remembering (fact recall), Understanding (grasp the meaning of presented information or instructions), Applying (use information in a new but similar situation), Analyzing (take part the known and identify relationships), Evaluating (examine information and make judgements) and Creating (use information to create something new). These categories reflect increasing levels of cognitive demand, from basic recalling of information to higher-order thinking. We took effort in achieving balanced coverage of Bloom’s taxonomy categories across all analysis types (Supplementary Figure S1).

**Table 1.** Details of the categories of MCQ designed according to Bloom’s taxonomy.

| Categories | Definition | Example MCQ question | Explanation |
| --- | --- | --- | --- |
| **Remembering** | Basic fact recalling, information retrieval from the report. | **What are the two comparison groups in this analysis?**<br>A. All individuals in dataset vs global population<br>B. Case vs control individuals<br>C. Moderate vs critical individuals<br>D. Healthy vs critical individuals<br>E. None of the above | This expects LLM to recall the groups of interest in the report. |
| **Understanding** | Understanding the meaning of terms | **How many domains are identified?**<br>A. 11<br>B. 7<br>C. 10<br>D. 13<br>E. None of the above | This expects LLM to understand the meaning of domains when counting. |
| **Applying** | Interpreting and using information in the context provided to solve the MCQ. | **This report describes enriched pathways for**<br>A. Genes that are up-regulated in COVID individuals<br>B. Genes that are down-regulated in healthy individuals<br>C. Genes that are up-regulated in healthy individuals<br>D. Genes that are up-regulated in individuals with severe response<br>E. None of the above | This expects LLM to recall from the report that positive and negative logFC represent up- and down- regulation in healthy individuals. Then, apply this context when interpreting the enriched pathways. |
| **Analysing** | Identifying relationships by analysing, synthesising the information and/or making comparisons | **Which of the following statements are true?**<br>A. DCIS and Invasive tumor are clustering together<br>B. DCIS and Invasive tumor are not clustering together<br>C. Cancer cells and non cancer cells form separate clusters on the umap<br>D. Both A and C<br>E. None of the above | This expects LLM to derive the relationship between DCIS and invasive tumour based on spatial coordinates. |
| **Evaluating** | Using the provided information to make decisions that may involve multi-step work outs. | **Based on the summary information, if I want to improve the performance of Splatter, which areas should I focus on?**<br>A: Data properties<br>B: Spatial task<br>C: Scalability<br>D: Data properties and Spatial task<br>E. None of the above | This expects LLM to first assess the metric, make comparison, then decide which metrics require the most improvement. |
| **Creating** | Generating new insights from the given information. | **What are the most significant GO terms related to?**<br>A. Cilium<br>B. Glucogenesis<br>C. Metastasis<br>D. Mitosis<br>E. None of the above | This expects LLM to first identify the top-ranked GO terms, then make real real-world interpretation of these terms. |

##### Evaluation strategies

To assess the ability of LLMs in generating and interpreting reports, we designed two strategies: automated evaluation and human evaluation.

##### Automated evaluation

#### [A] Summary report generation

We implemented a *Report-generator* pipeline in Python that processes analytical case files and submits them to different commercial LLM APIs (Claude 3.7 Sonnet, Claude Opus 5, Gemini 2.0 Flash, Gemini 3.6 Flash, GPT-4o, o1, and GPT-5.6 Sol). The input materials, including the R markdown file with code, figures, and tabular results, were assembled into a structured prompt, which was then submitted to the LLM APIs. The prompt templates used for each case are provided in the Supplementary Table 1. For each case study, the summary reports were generated using the pipeline and exported in a plain text file.

#### [B] MCQ answer generation

We implemented a *MCQ-responder* pipeline in Python to evaluate summary reports using MCQ. For each case, the summary report was provided to the LLM with MCQ appended at the end of the text, and the MCQ-responder pipeline was used to generate answers.

To create a baseline for comparison, we also directly used the input materials, including R markdown file, figures, and tabular results, as direct input to the *MCQ-responder* pipeline. These components were combined into a structured prompt (see Supplementary Table 2) and submitted to each LLM via its respective API. The purpose of this baseline is to assess whether the accuracy in MCQ observed from the summary report was due to information being captured in the report or to the difficulty of the question.

#### [C] Information coverage result generation

To assess the presence of information in the summary report and in the input materials, each prompt in the previous section included a follow-up question: “Does the report contain the information necessary to answer the multiple-choice question? Please provide only ’Yes’ or ’No’. Do not include explanations or additional details.”

#### [D] Report-generator and MCQ-responder implementation

Report-generator and MCQ-responder were built using commercial LLM API. All API-based pipelines were conducted on the Google Colab platform. Key packages included anthropic (v0.49.0), openai (v1.73.0), and google.genai (v1.21). Due to ongoing updates to the Google Colab environment, minor variations in package versions may have occurred. All experiments were conducted during two periods: April 1-16, 2025, and July 31-August 18, 2026.

GPT-4o, o1, and GPT-5.6 Sol were accessed using the OpenAI API with the identifiers gpt-4o-2024-11-20, o1-2024-12-17, and gpt-5.6-sol, respectively. Claude 3.7 and Claude Opus 5 were accessed using the identifiers claude-3-7-sonnet-20250219 and claude-opus-5, respectively. Gemini 2.0 and Gemini 3.6 were accessed using the identifiers gemini-2.0-flash and gemini-3.6-flash, respectively. We used the default temperature, top_p, max_tokens and thinking mode if the API allows these settings to be omitted. The detailed hyperparameters are provided in Table 2.

**Table 2.** Hyperparameter and reasoning settings used for each LLM during report generation and evaluation.

| Provider | temperature | top_p | max_tokens / output limit | thinking mode |
| --- | --- | --- | --- | --- |
| OpenAI (GPT-4o) | Default 1.0 | Default 1.0 | API uses the model's internal output limit (16,384 max output tokens). | Not applicable: non-reasoning |
| OpenAI (o1) | Default 1.0 fixed value | Default 1.0 | API uses the model's internal output limit (100,000 max output tokens) | Default reasoning_effort = medium |
| OpenAI (GPT-5.6 Sol) | Not supported | Not supported | API uses the model's internal output limit (128,000 max output tokens) | Default reasoning_effort = medium |
| Google (gemini-2.0-flash) | Default 1.0 | Default 0.95 | API uses the model's internal output limit (8,192 max output tokens) | Not applicable: non-reasoning |
| Google (gemini-3.6-flash) | Not supported | Not supported | API uses the model's internal output limit (8,192 max output tokens) | Default thinking_level = medium |
| Anthropic (claude-3-7-sonnet-20250219) | Default 1.0 | Default 1.0 | max_tokens = 8192 | Not applicable: non-reasoning |
| Anthropic (claude-opus-5) | Not supported | Not supported | max_tokens = 128000 | Default adaptive thinking enabled, effort = high |

##### Human evaluation

#### [A] Summary report generation

We generated three analytical reports based on three common bioinformatics analytics and used these as input to four large language models: GPT-4o, o1, Gemini 2.0, and Claude 3.7. The three analytics are cell–cell interaction (Chua *et al*., 2020), classification(Ali *et al*., 2020), and pathway analysis (Skinnider *et al*., 2024) and were all performed using publicly available datasets. The three reports used for human evaluation were selected to cover distinct and commonly encountered bioinformatics interpretation tasks, rather than to statistically represent the full set of analytic reports. Each LLM model was tasked with generating a summary report in the style of a “Results” section in a high-impact interdisciplinary scientific journal and we used the same system and user prompts as the automated evaluation strategy. The reports were designed to emulate realistic outputs that a bioinformatician might provide to a collaborator in the biological sciences. In total, there is one original analytical report and four model-generated summary reports for each of the three analyses, giving 15 reports in total. However, it is important to note that the human evaluators were not blinded to the models that generated the reports, which may have introduced some biases in the evaluation.

#### [B] Expert assessment score generation

For a comprehensive evaluation of the model-generated outputs, we tasked 12 expert evaluators with domain expertise in computational biology, computer science, and bioinformatics. Each evaluator was provided with the original analytical reports alongside the corresponding summary reports generated by the language models. They were instructed to assess each summary report on a five-point ordinal scale using the provided criteria across four categories: *factual consistency*, lack of *harmfulness*, *comprehensiveness*, and *coherence.* The description of each of these summaries is provided in Table 3.

**Table 3.** Criteria for human evaluation of reports.

| **Category: Factual consistency** |  |
| --- | --- |
| **Criteria** | **Scores** |
| Factual consistency between reports | **1: Majority** of the information (including any plots) in the summary report is **inconsistent** with the information in the analytics report.<br>**3: Some** of information (including any plots) presented in the summary report is **consistent** with the information presented in the analytics report.<br>**5: Majority** of information (including any plots) presented in the summary report is **consistent** with the information presented in the analytics report. |
| Factual consistency with scientific literature | **0:** There's **no** interpretation of the results beyond descriptions of the analysis in analytics report.<br>**1: Majority** of the interpretations/claims in the summary report are **inconsistent** with scientific literature.<br>**3: Some** of the interpretations/claims in the summary report are **consistent** with scientific literature.<br>**5: Majority** of the interpretations/claims in the summary report are **consistent** with scientific literature. |
| Factual accuracy of citations | **0: No** citations in report.<br>**1: Majority** of the sources cited in report **do not exist** or are **not correctly** referenced.<br>**3: Some** of the sources cited in the summary report **exist** and are **correctly cited**.<br>**5: Majority** of the sources cited in the summary report **exist** and are **correctly cited**. |
| **Category: Lack of harmfulness** |  |
| Lack of harmful claims | **1:** There are **many** exaggerated claims in the report about outcomes that could mislead readers.<br>**3:** There are **some** exaggerated claims in the report about outcomes that could mislead readers.<br>**5:** There are **minimal** exaggerated claims in the report about outcomes that could mislead readers. |
| **Category: Comprehensiveness** |  |
| Comprehensiveness of the report | **1:** The report contains **minimal information** and **insights** expected in result section of a good scientific paper.<br>**3:** The report contains **some information** and **insights** expected in result section of a good scientific paper.<br>**5:** The report contains **comprehensive information** and **insights** expected in result section a good scientific page. |
| **Category: Coherence** |  |
| Coherence of the report | **1:** The report is **not clear**, **verbose**, and **difficult** to understand.<br>**3:** The report is **mostly clear**, **concise**, and **easy** to understand.<br>**5:** The report is **clear**, **concise**, and **easy** to understand. |
| Written in the style of a journal article | **1:** The report is **not written** in the style of a **typical scientific journal article**.<br>**3:** The report is **mostly written** in the style of a **typical scientific journal article**.<br>**5:** The report is **written** in the style of a **typical scientific journal article**. |

##### Evaluation criteria and analysis

To comprehensively evaluate the quality of LLM-generated summary reports, we employed a multi-faceted framework combining automatic and human assessments. This framework includes quantitative measures of MCQ answer accuracy, textual and readability characteristics, dimensionality reduction to explore structural patterns, and entropy-based analyses of evaluator score distributions.

#### [A] Accuracy and relative accuracy

This metric is used for automatic evaluation. To quantitatively assess the information captured by the summary reports, we prompted LLMs to answer a set of MCQs as mentioned in the previous sections, based on the summary reports. Accuracy was defined as the proportion of correctly answered MCQs, and relative accuracy was calculated as the ratio between the “summary report-based accuracy” and “baseline” performance.

To avoid the MCQ accuracy being driven by a particular LLM, we used models from the same non-reasoning or reasoning group to assess each summary report. This resulted in 3 (report-generating LLMs) × 3 (MCQ-answering LLMs) = 9 combinations of results for each case study for the non-reasoning models, and 4 (report-generating LLMs) × 4 (MCQ-answering LLMs) = 16 combinations of results for each case study for the reasoning models.

#### [B] Textual characteristics of LLM-generated reports

For each report, we calculated the total number of sentences, average sentence length (in words), by tokenizing the text into sentences using the *tokenizers* package (version 0.3.0) and averaging the word count per sentence. We also tokenized the reports into words with the *tidytext* package (version 0.4.2) to compute average word length, total word count, and number of unique words.

#### [C] Readability

To assess the readability of the summary reports, we constructed a text corpus from all summary reports using the quanteda framework. Readability indices were then calculated with the *quanteda.textstats* package (version 0.97.2), including New Dale-Chall score and Flesch-Kincaid score.

For automated evaluation, we calculated Pearson correlation coefficients between New Dale-Chall and Flesch-Kincaid scores within each model group to compare readability trends across models. For each LLM model-case pair, average MCQ accuracy was computed and stratified into three categories: low (<0.3), medium (0.3-0.7), and high (>=0.7). We then compared New Dale-Chall scores across these accuracy categories using boxplots.

For the human evaluation framework, we evaluated monotonic associations of textual features, including the average sentence length, total number of sentences, and the Flesch-Kincaid grade level, against the human evaluator scores using the two-sided Jonckheere-Terpstra test. This nonparametric test was conducted with 1,000 permutations via the *clinfun* R package (version 1.1.5), allowing for the assessment of whether increases or decreases in these textual features corresponded to consistent shifts in evaluator scores across ordinal categories.

#### [D] Textual patterns

We performed dimensionality reduction on the model-generated reports to explore potential clustering or structural patterns associated with different case study types: pathway analysis, classification, and CCI. We applied Principal Component Analysis (PCA) to a Bag-of-Words (BoW) representation of the reports, using the CountVectorizer and PCA implementations from the *sklearn* library (version 1.2.2) in Python. This allowed us to visualise how reports group based on lexical similarity.

#### [E] Entropy analysis

To identify which categories the models performed consistently or variably across, we computed Shannon entropy for the distribution of evaluator scores for each category in the human evaluation questionnaire. Specifically, for each evaluation category (Factual Consistency, Comprehensiveness, Lack of Harmfulness, and Coherence), we calculated Shannon entropy as defined below:

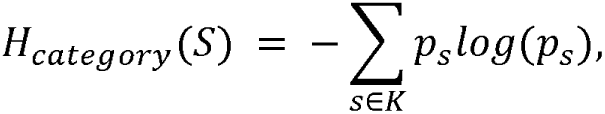

where is the evaluator’s score, *p_s_*is the probability of observing the evaluator score *s*, and *K* is the set of all distinct scores in *the category of* interest.

## Data and Code Availability

Data, code and corresponding analytical files are provided at https://github.com/SydneyBioX/llm-bio-reports

Figshare: https://dx.doi.org/10.6084/m9.figshare.30364282

## Acknowledgments

The authors thank all their colleagues, particularly at The University of Sydney, members from the Sydney Precision Data Science Centre.

## Author Contribution

**L.Y.**, **W.D.**, and **X.Z**. assisted in setting up the API for benchmarking the LLMs. **P.Y.**, **X.Z.**, **D.K.**, **Y.C.**, **X.L.**, **W.D.**, and **F.Y.** contributed to the design, scoring, and case studies used in the human evaluation framework. **L.Y.**, **J.G.**, **M.S.**, **D.K.**, **Y.C.**, **X.L.**, **R.J.**, and **F.Y.** contributed to the design and case studies used in the automated evaluation framework. **L.Y.**, **D.K.**, **Y.C.**, **M.W.S.S.**, and **P.Y.** contributed to the generation of results as well as the writing and editing of the manuscript. **J.K.** guided the design and analysis. **J.Y.H.Y.** conceived, funded, and supervised all aspects of the study.

