## Supplemental Figures for "Large Language Models for Accessible Reporting of Bioinformatics Analyses in Interdisciplinary Contexts"

### Supplementary Methods

#### [A] Prompt engineering

For all LLMs, that is, the report generating LLM and MCQ answering LLM in strategy 1 and the MCQ answering LLM in strategy 2, we used a two-part prompt design that includes a system prompt and a user prompt. The system prompt instructs the LLM to act as an expert scientist with a strong background in bioinformatics and biology. It also provides the instruction about its task. For the report generating LLM, its task is to summarise the analytical report into a logical narrative. For the MCQ answering LLM, its task is to answer the multiple choice based on the summary report and output a single option only. The system prompt is followed by the user prompt that informs LLM about the files being uploaded.

#### [B] Supplementary table

**Supplementary Table 1. Prompts used for the automated evaluation framework**

| **Automated evaluation framework** | |
| --- | --- |
| **Report generating LLM (System prompt)** | |
| Old | New |
| You are a highly skilled expert biological writer tasked with generating a result paragraph in a high profile scientific journal such as Science, Nature, or Cell in a concise format. | I am a bioinformatician that has conducted some analyses but lacks the biological knowledge to interpret the results and build a cohesive narrative. However, you are an expert scientist who has a strong background in both bioinformatics and biology. I would like to ask you to interpret and translate my bioinformatics analyses into a coherent “Results” section suitable for a high-impact interdisciplinary scientific journal.  I have provided you with a .txt file containing the outputs from my analysis, and in some cases, additional supporting materials such as plots. These files are the core analytics you will need to synthesize and summarize into a coherent narrative. The .txt file also includes an “LLM” detailing any specific requirements or instructions for your summary. Please read it carefully.  Please remember the following when completing this request: Interpret and synthesize the provided bioinformatics results into a clear and logical narrative, complete any specific requests outlined in the LLM task, include quantitative evidence from the analyses to support your conclusions, and include relevant literature to support or contextualise your findings in your summary. |
| **Report generating LLM (User prompt)** | |
| Old | New |
| I have a rmd text file: I have multiple csv files: I have image files:  Please summarize the content of all the files above and the attached images to answer the LLM task described at the beginning of the rmd text file, ensuring that as many details as possible are preserved. | I have provided you with a .txt file containing the outputs from my analysis, and in some cases, additional supporting materials such as plots. Please synthesize and summarize the contents of the file(s) into a coherent narrative. Be sure to read the “LLM task” in the .txt file.   Txt file: CSV file: Image file: |
| **MCQ answering LLM (System prompt)** | |
| Old | New |
| You must respond concisely by selecting only the appropriate option from A, B, C, D, or E when applicable. For yes/no questions, respond strictly with 'Yes' or 'No.' Do not provide any explanations, commentary, or additional text beyond the selected option. | You are an expert scientist who has a strong background in both bioinformatics and biology. I have provided you with a .txt file containing the outputs from my analysis, and in some cases, additional supporting materials such as plots. These files are the core analytics you will need to understand to read the following multiple choice question. When answering the multiple choice question, please select one option only. Please be concise without including any additional information. For example if you think option A is the correct answer, then please just output A. |
| **MCQ answering LLM (User prompt)** | |
| Old | New |
| This is a report: XXXX From the report, I want to know: MCQ XXXXXX  Please provide only the letter of the correct option (A, B, C, D, or E). Do not include the answer text, explanations, or any other information. | Please read the following report and use the information to answer the multiple choice question below.  Report: XXX  Multiple choice question: XXX  Please provide only the letter of the correct option (A, B, C, D, or E). Do not include the answer text, explanations, or any other information. |
| Does the report contain the information necessary to answer the multiple-choice question? Please provide only 'Yes' or 'No'. Do not include explanations or additional details. | Does the report contain the information necessary to answer the multiple-choice question? Please provide only 'Yes' or 'No'. Do not include explanations or additional details. |

**Supplementary Table 2. Prompts used for sensitivity analysis baseline**

| **Sensitivity analysis baseline** | |
| --- | --- |
| **MCQ answering LLM (System prompt)** | |
| Old | New |
| You are a highly skilled scientist with expertise in synthesizing complex scientific information. You will receive multiple files containing relevant data along with a single multiple-choice question. Your task is to integrate and summarize the provided content to accurately answer the question. Your response must consist solely of the letter corresponding to your chosen answer (A, B, C, D or E) with no additional commentary. | You are an expert scientist who has a strong background in both bioinformatics and biology. I have provided you with a .txt file containing the outputs from my analysis, and in some cases, additional supporting materials such as plots. These files are the core analytics you will need to understand to read the following multiple choice question. When answering the multiple choice question, please select one option only. Please be concise without including any additional information. For example, if you think option A is the correct answer, then please just output A. |
| **MCQ answering LLM (User prompt)** | |
| Old | New |
| Please summarize the content of all the files above and the attached images based on the LLM task described at the beginning of the rmd text file, and answer the following question: XXXXX. Please provide only the letter of the correct option (A, B, C, D, or E). Do not include the answer text, explanations, or any other information. | Please read the following .txt file and any supporting materials and use the information to answer the multiple choice question below.  Please provide only the letter of the correct option (A, B, C, D, or E). Do not include the answer text, explanations, or any other information.  Txt file: XXX CSV file: XXX Image file: XXX  Multiple choice question: XXX |
|  | Please indicate whether the information needed to answer the multiple-choice question can be found in the document. Please provide only 'Yes' or 'No'. Do not include explanations or additional details. |

### **Supplementary Figures**

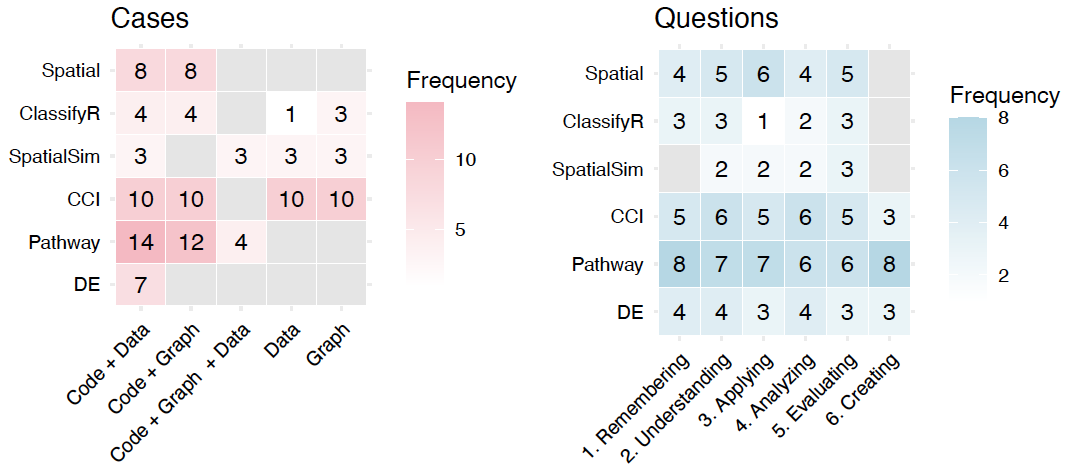
**Supplementary Figure S1.Number of cases and questions across input type and information retrieval strategy.** In total, we generated 117 cases and 138 multiple choice questions.

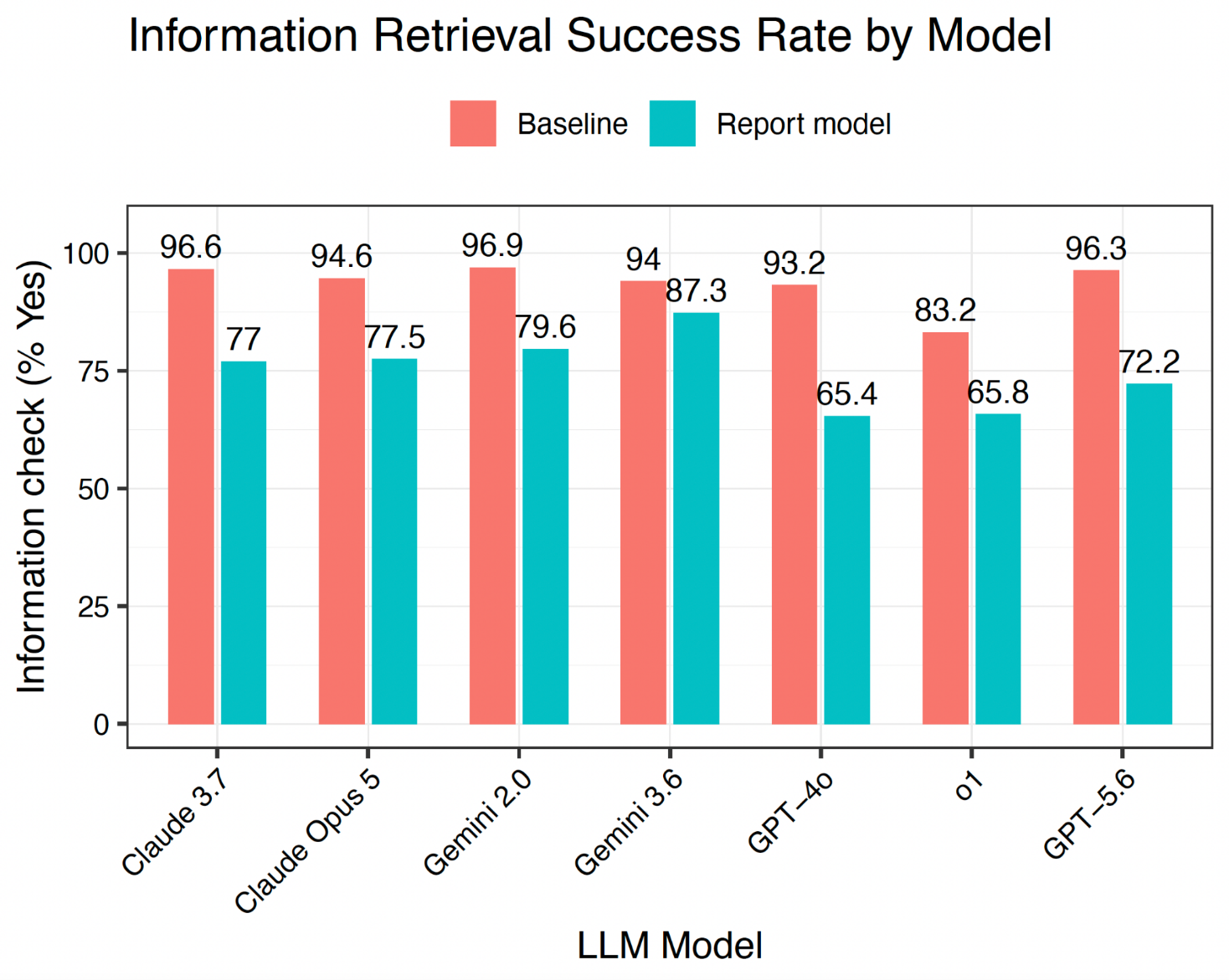

**Supplementary Figure S2. Information coverage comparison across the baseline model and report model.** The percentage is calculated by averaging across reports generated by each report-generator LLM.

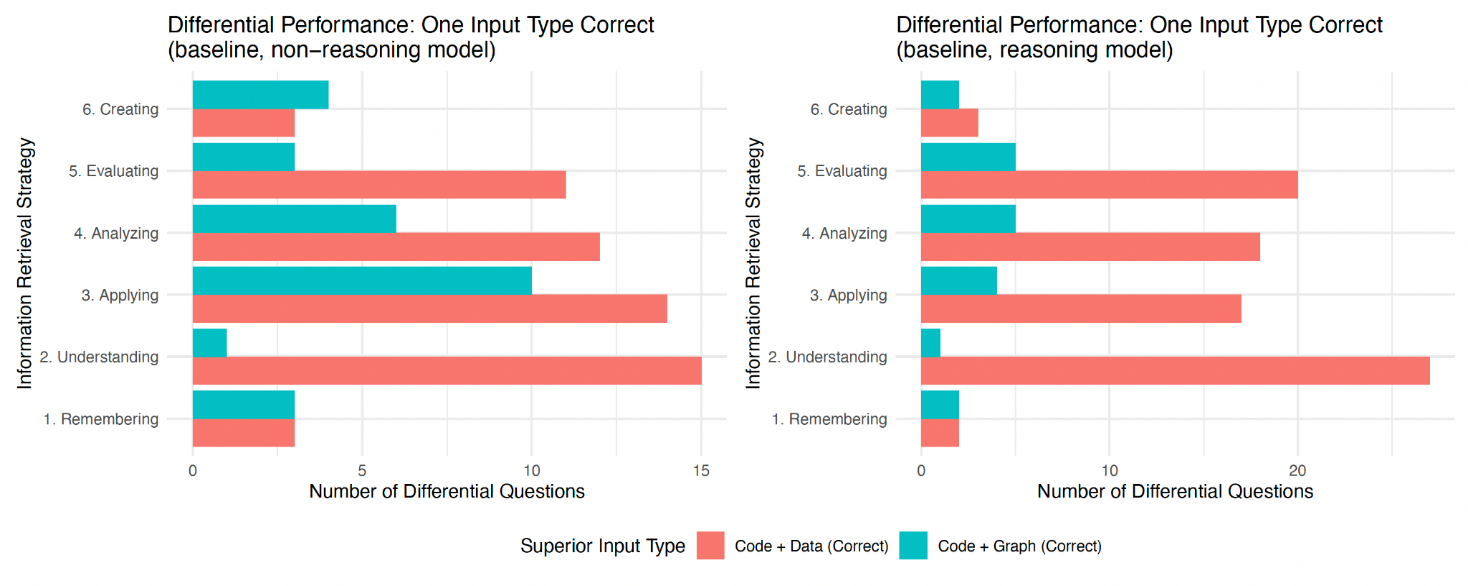

**Supplementary Figure S3: MCQs where a certain input type was correct.**

We examined MCQs that were answered correctly by either Code + Data or Code + Graph but not both. The bar indicates the number of MCQs where a certain input type was correct, stratified by the type of MCQs.

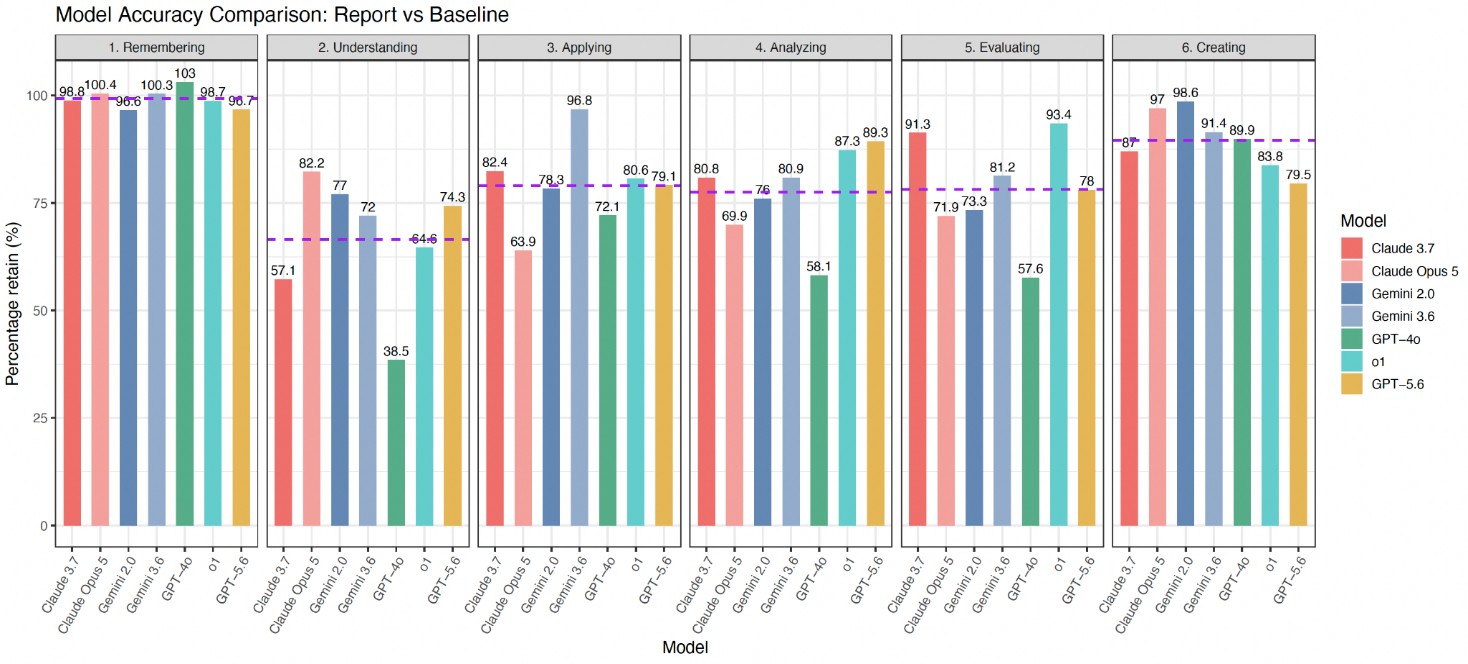

**Supplementary Figure S4: Percentage of information retention across information retrieval strategy.** We examined the percentage of information retention in the report model and in the baseline line model. Percentage retained is calculated as the ratio of the information retention in the report models and in the baseline models.

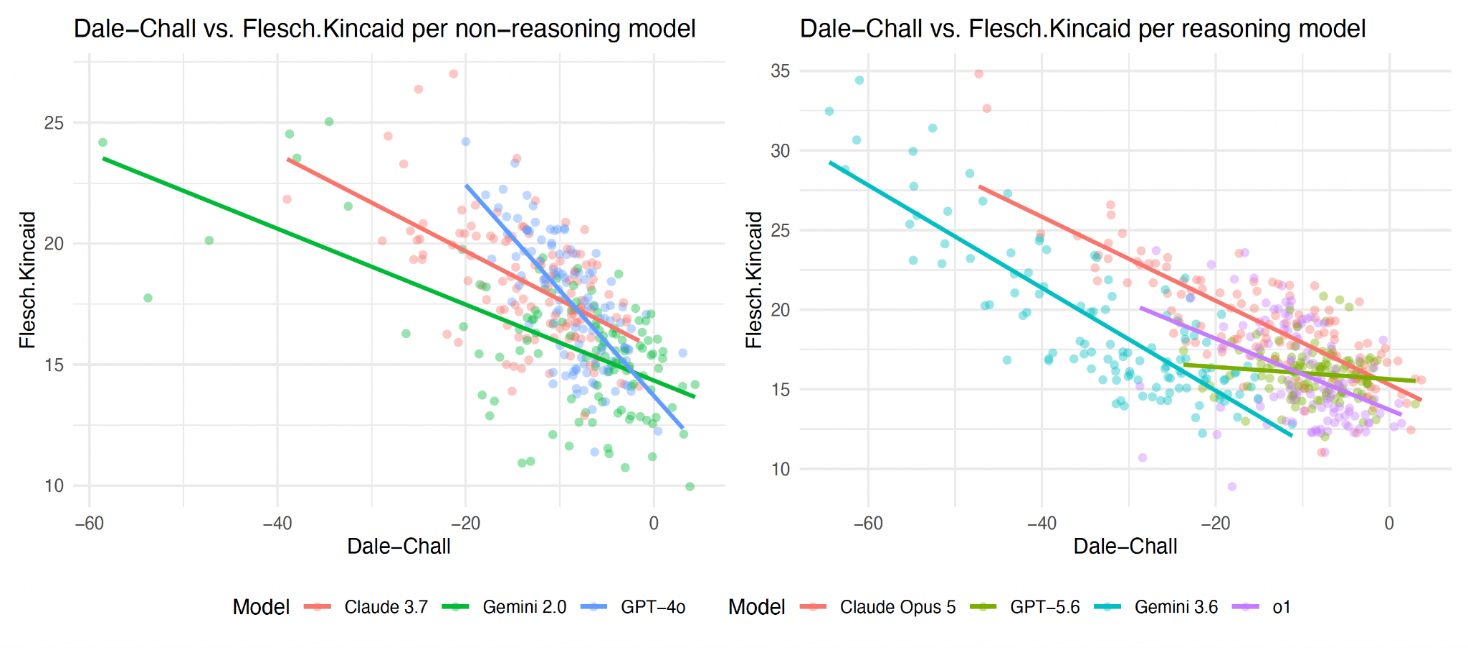

**Supplementary Figure S5: Dale-Chall and Flesch Kincaid value for each LLM generated report.** Each dot on the plot represents a report with a linear regression line fitted to the reports generated by each report-generating LLM model.

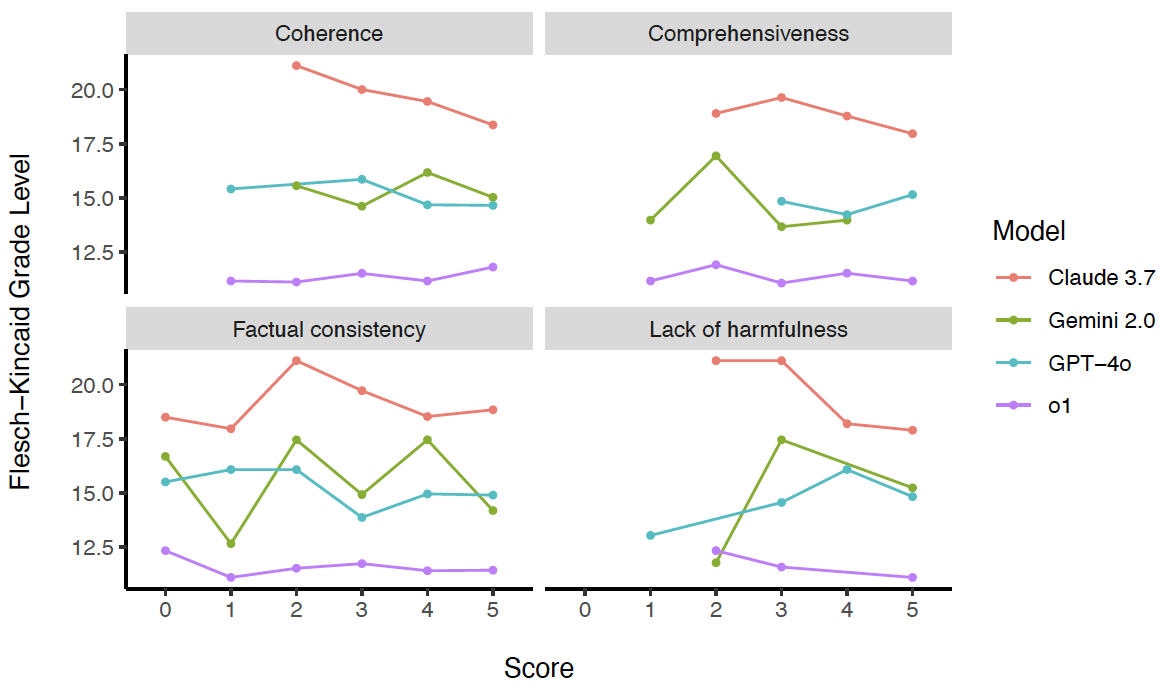

**Supplementary Figure S6**. Flesch-Kincaid Grade Levels for Human evaluation framework.

The Flesch-Kincaid Grade Level was computed for all models across the four categories defined in the human evaluation framework. Higher scores reflect more complex, less accessible language, indicating that a higher school grade level would be required to understand the output.

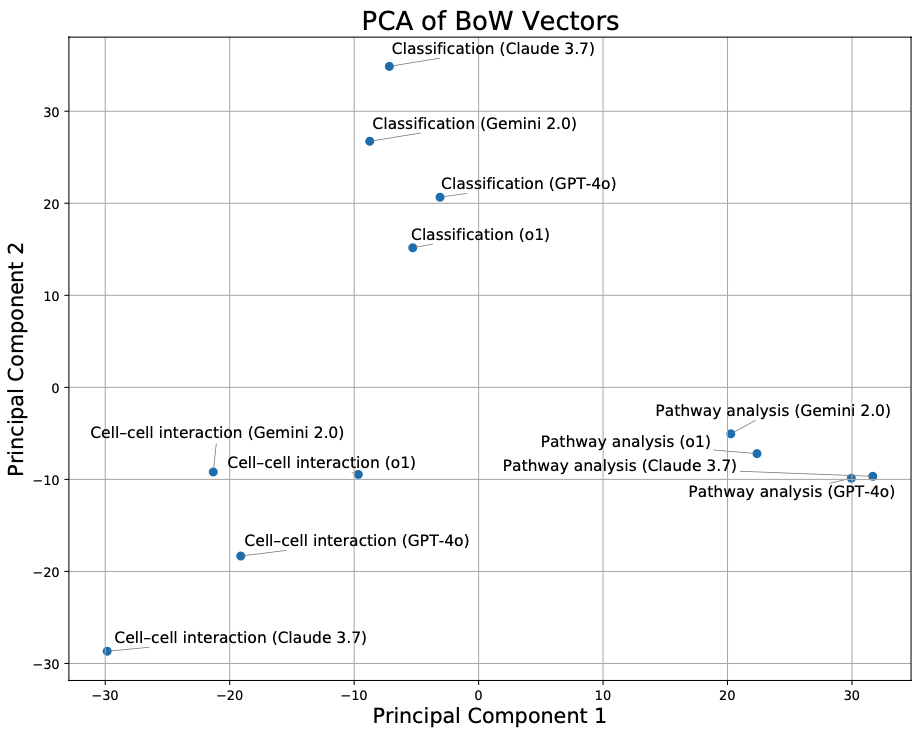

#### Supplementary Figure S7. PCA of reports assessed in manual evaluation. We used the Bag-of-Words representation to convert reports into features for PCA. Each point represents a report of a particular analysis (Pathway analysis, Cell-cell interaction, Classification) output by a particular report generator LLM (Gemini 2.0, Claude 3.7, GPT-4o, o1).

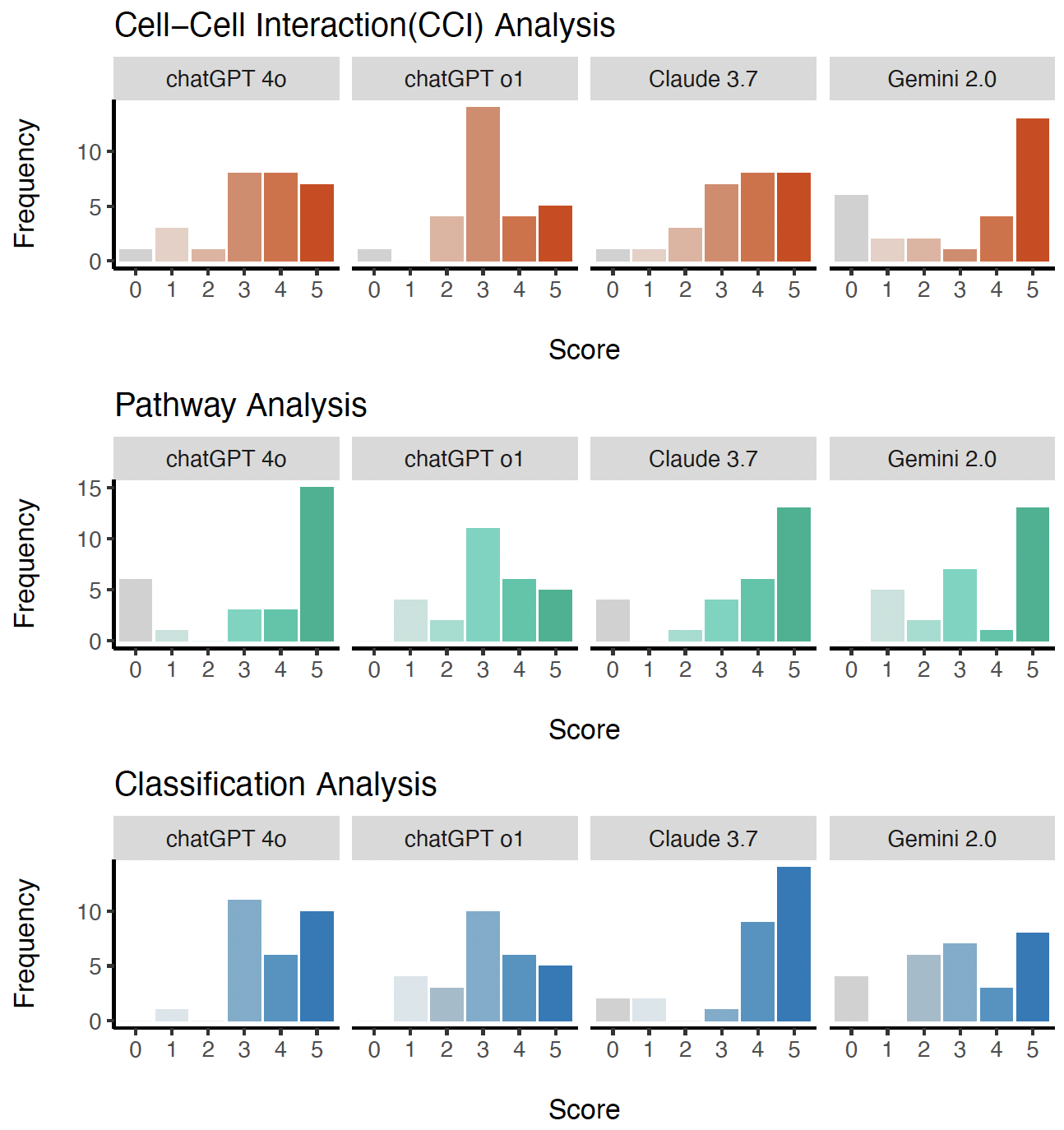

#### Supplementary Figure S8. Scores from human assessors on the reports selected for manual evaluation. In manual evaluation, every report is assessed from multiple criteria of factual consistency, coherence, comprehensiveness and lack of harmfulness, where each question is graded from 0 to 5. The barplot summarises the scores received across all criteria for each report.

**Supplementary File S9**

Supplementary File S9 presents the exact user prompts used across three prompt-engineering iterations and the corresponding large language model (LLM) outputs when tasked with translating bioinformatics analysis reports into a journal-style Results section. The prompts progressively introduce (i) clearer task instructions, (ii) explicit requirements to include quantitative evidence, and (iii) additional contextual framing about the intended audience and the model’s assumed expertise. The resulting outputs demonstrate how prompt specificity and contextual information substantially influence the coherence, relevance, and evidentiary rigor of generated summaries. This file serves as an empirical record of the prompt design process underpinning the benchmarking and evaluation strategies described in the manuscript.

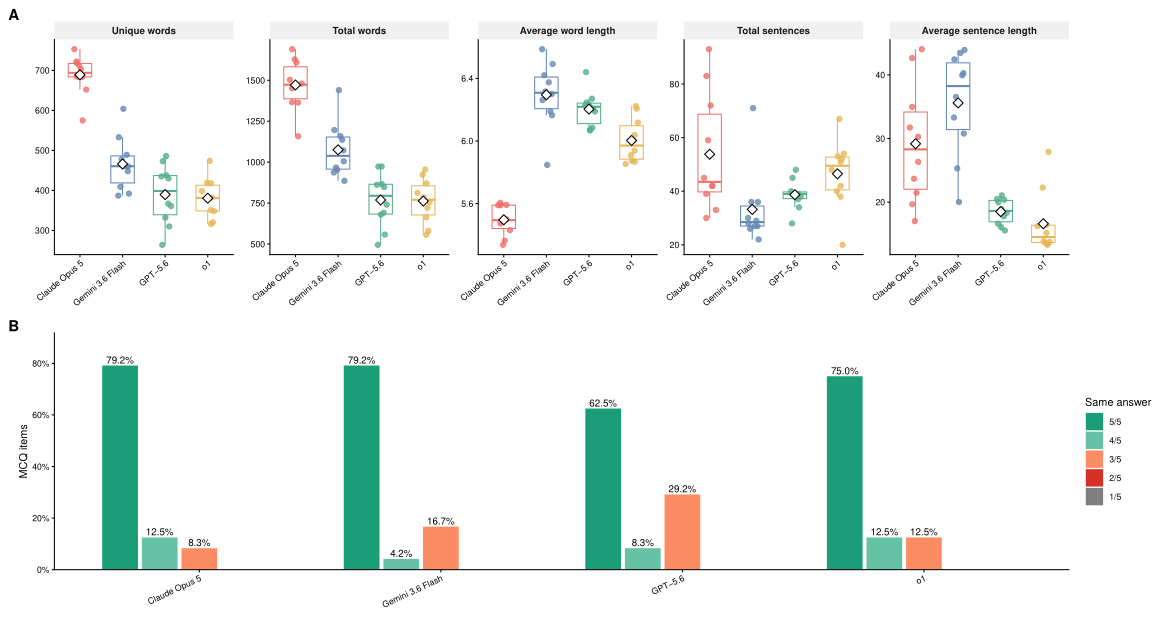

**Supplementary S10 Repeated-run sensitivity analysis of LLM-generated reports and automated evaluation.** (A) Variation in report characteristics across five repeated generations for four reasoning models using the Kidney pathway 1 case study. Diamonds indicate mean values. (B) Consistency of automated MCQ answers across five repeated evaluation runs. Bars show the percentage of MCQ items receiving the same answer in 1-5 runs.
